# Natrix: A Snakemake-based workflow for processing, clustering, and taxonomically assigning amplicon sequencing reads

**DOI:** 10.1101/2020.09.23.309864

**Authors:** Marius Welzel, Anja Lange, Dominik Heider, Michael Schwarz, Bernd Freisleben, Manfred Jensen, Jens Boenigk, Daniela Beisser

## Abstract

Sequencing of marker genes amplified from environmental samples, known as amplicon sequencing, allows us to resolve some of the hidden diversity and elucidate evolutionary relationships and ecological processes among complex microbial communities. The analysis of large numbers of samples at high sequencing depths generated by high throughput sequencing technologies requires efficient, flexible, and reproducible bioinformatics pipelines. Only a few existing workflows can be run in a user-friendly, scalable, and reproducible manner on different computing devices using an efficient workflow management system. We present Natrix, an open-source bioinformatics workflow for preprocessing raw amplicon sequencing data. The workflow contains all analysis steps from quality assessment, read assembly, dereplication, chimera detection, split-sample merging, sequence representative assignment (OTUs or ASVs) to the taxonomic assignment of sequence representatives. The workflow is written using Snakemake, a workflow management engine for developing data analysis workflows. In addition, Conda is used for version control. Thus, Snakemake ensures reproducibility and Conda offers version control of the utilized programs. The encapsulation of rules and their dependencies support hassle-free sharing of rules between workflows and easy adaptation and extension of existing workflows. Natrix is freely available on GitHub (https://github.com/MW55/Natrix).

## 1 Background

Prokaryotes and microbial eukaryotes constitute a large fraction of the biodiversity on earth, but in many environments, their distribution and diversity are still unknown [1]. Sequencing of marker genes amplified from environmental samples can resolve some of the hidden diversity and elucidate evolutionary relationships and ecological processes among the microorganisms. In particular, the introduction of high-throughput technologies and sequencing of genomic DNA regions (such as the 16S or 18S rRNA genes) and the internally transcribed spacer region at high sequencing depths enable detailed analyses and profiling of complex microbial communities [2]. The analysis of large sample numbers at high sequencing depths generated by recent Illumina sequencing technology requires efficient, flexible, and reproducible bioinformatics workflows. There are several existing bioinformatics tools available, including QIIME2 [3], USEARCH [4], Fred’s metabarcoding pipeline [5], and mothur [6]. However, only a few tools include all necessary analytic steps starting from raw sequencing reads up to taxonomically assigned sequence representatives and can be used to execute user-friendly, scalable, and reproducible workflows on different computing devices. Existing tools often contain a collection of bash scripts that are run successively on each sample without using an efficient workflow management system.

In this paper, we present Natrix, an open-source bioinformatics workflow that allows researchers without programming experience to process their amplicon data using a single command, while being easy to modify by advanced users. The workflow is written using Snakemake [7], a workflow management engine for developing data analysis workflows. Snakemake ensures reproducibility of a workflow by automatically deploying dependencies of workflow steps (rules) and scales seamlessly to different computing environments like servers, computer clusters, or cloud services. Furthermore, Conda (https://docs.conda.io/en/latest/), and specifically Bioconda [8], are used for version control of the utilized programs, leading to a simple installation without the risk of dependency conflicts. The workflow contains separate rules for each step and each rule that has additional dependencies has a separate Conda environment that will be automatically created when starting the workflow for the first time. The encapsulation of rules and their dependencies supports hassle-free sharing of rules between workflows and easy adaptation and extension of existing workflows. We briefly compare Natrix to existing pipelines for amplicon analysis and evaluate the features, advantages, and disadvantages of each approach.

## 2 Methodology and Implementation

### Overview

Natrix has two main workflow variations depending on the chosen sequence representation (Fig. 1): one for amplicon sequence variants (ASVs), using DADA2 [9] for inference of sequencing variants, and one for operational taxonomic units (OTUs) based on the swarm clustering algorithm [10]. The main difference between the two variants is during which point of the workflow the sequence representatives are inferred. Natrix is highly customizable and easy to use without programming experience. Out of the box, it supports Illumina single-end and paired-end data and the AmpliconDuo split-sample protocol developed by [11]. Many of the workflow steps are optional and can be deactivated in the configuration file. The user can also choose the reference database for the taxonomic assessment. Currently, SILVA [12] and NCBI [13] can be used without modifications, while alternative databases can easily be integrated. Natrix is written in Snakemake [7], a workflow management engine inspired by GNU Make using a Python-like language. The generation of output files from input files in Snakemake is defined in a rule. The definition of a rule generally specifies the name of the rule, one or more input and output files and a shell command or the path to a Python or R script that creates the output file(s) from the input file(s). It can optionally contain additional parameters, a path to a Conda environment (https://docs.conda.io/en/latest/) to deploy dependencies of workflow steps, paths to log files and benchmark files, and restrictions of the resources (like the number of threads or the amount of memory) that can be used by a rule. The order in which the rules are executed and which rules can be run in parallel is automatically inferred by Snakemake when a workflow is started by computing a directed acyclic graph (DAG). If a target file is not specified during the start of the workflow, Snakemake will infer the order based on the input required by the first rule, which creates an *all* rule that exclusively contains the desired target files of the workflow as input. The main workflow is configured using a configuration file (additional file 3), where each option is documented. During the initialization of a workflow process, all values of the configuration file are validated for logical correctness, reducing the probability of faulty workflow outputs due to typographical errors. On a local computer and on a remote server, a workflow can be started out of the box with a single command. Using Natrix in distributed computing environments requires minimal adjustment for the software stack available in the environment.

**Figure 1:**
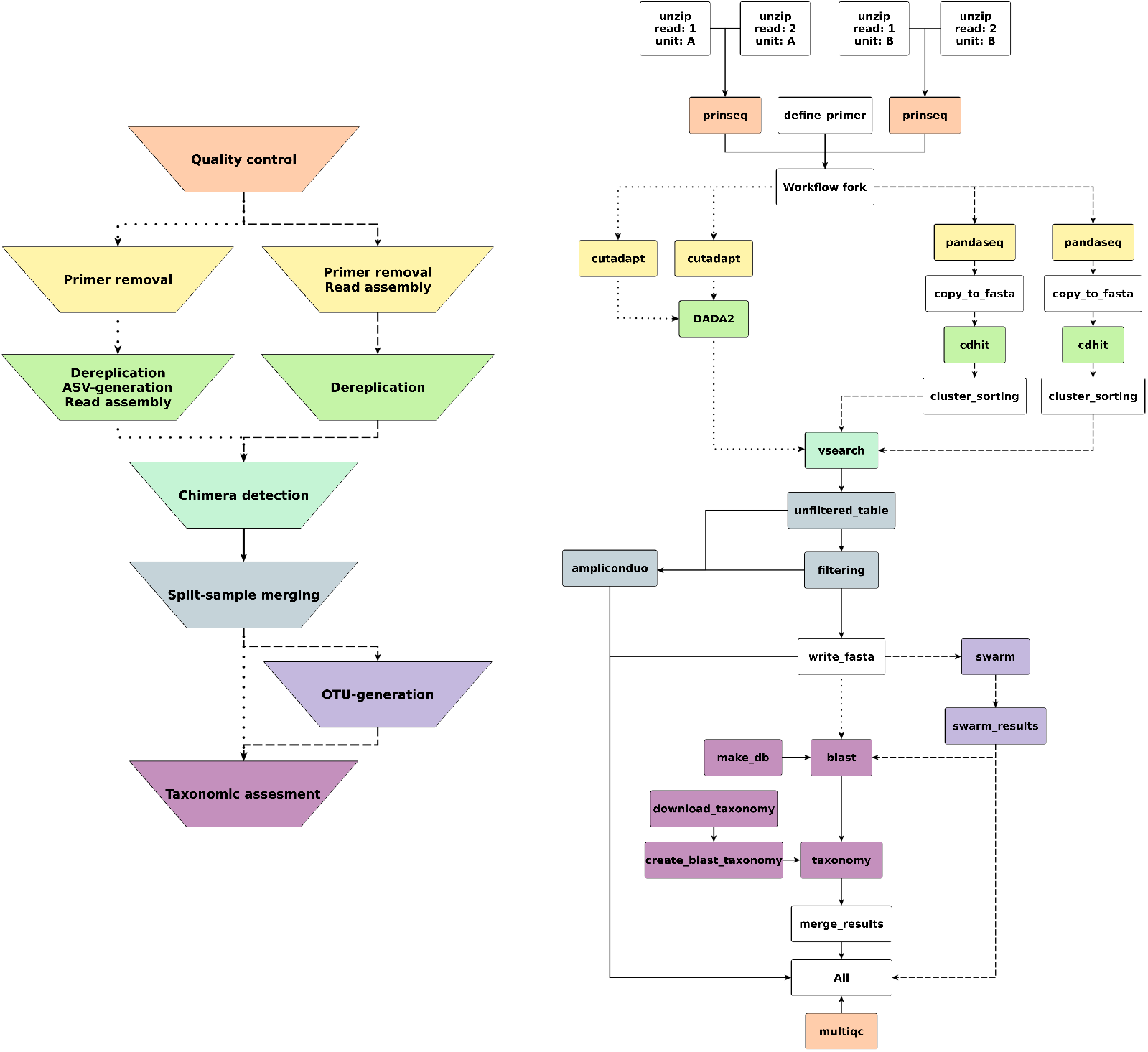
Left: schematic representation of the main steps of the workflow; the color coding represents which rules belong to which main step. Dotted edges denote the ordering of steps taken for the ASV variation of the workflow, dashed lines denote the ordering of steps taken by the OTU variation of the workflow, and straight lines are the steps that are identical in both versions. Right: Graph of an example workflow. Each node represents a rule instance to be executed. The direction of each edge represents the order in which the rules are executed. Disjoint paths in the graph can be executed in parallel.

### OTUs and ASVs

In 16S and 18S rRNA gene studies it is common to cluster sequences into OTUs. *De-novo* OTU generation methods often use an arbitrary clustering threshold (typically <97% sequence similarity) to cluster sequences [14]. By clustering similar sequences and taking the most abundant sequence as the representative of the cluster, the impact of sequencing or PCR errors can be reduced. Another reason for clustering sequences into OTUs is to account for intra-specific genetic diversity, allowing the use of OTUs as a proxy for species. One downside of this approach is that OTUs are not consistent between datasets, limiting comparability. Another disadvantage is the use of an arbitrary clustering threshold, but this can be alleviated with alternative clustering algorithms such as swarm [10], which uses an iterative approach of single-nucleotide differences between iterations to cluster sequences into OTUs. In recent years, another approach was developed for clustering Illumina sequences. This approach resolves ASVs without using arbitrary clustering thresholds and with increased resolution (up to a difference of a single nucleotide between ASVs) [9]. While there are arguments for completely abandoning OTUs in favor of ASVs [15], both serve different niches and should be chosen depending on the research question that is addressed. ASVs are consistent labels, allowing comparisons between datasets, and the increased resolution allows studying the distribution of gene polymorphisms over different datasets. Since OTUs serve as a proxy for species, they are well suited for studying the alpha-diversity of a dataset without having to account for intra-specific diversity in downstream analysis.

Since the different forms of sequence representation serve different niches, Natrix supports both the creation of ASVs using the DADA2 algorithm [9] and the picking of OTUs using the swarm clustering algorithm [10].

### Workflow steps

#### Initial demultiplexing

The demultiplexing step, i.e., sorting reads according to their barcode, is implemented in a separate script, independent of the rest of the workflow, since it is often already performed by the sequencing company.

#### Preprocessing

For quality control, the pipeline uses the programs FastQC [16], MultiQC [17], and PRINSEQ [18]. FastQC generates a quality report for each FASTQ file, containing information, such as the sequence quality per base and on the average (using the Phred quality score), overrepresented sequences, GC content, adapter, and the k-mer content of the FASTQ file. MultiQC aggregates the FastQC reports for a given set of FASTQ files into a single report, allowing reviews of all FASTQ files at once. PRINSEQ is used to filter out sequences with an average quality score below a threshold that can be defined in the configuration file of the workflow.

#### Read assembly, dereplication and removal of undesired subsequences

The subsequences that are specified by a primer table are removed before the assembly of forward and reverse reads and the dereplication of the dataset. A primer table contains information about the primer and barcode sequences used and the lengths of the poly-N subsequences. Besides removing the subsequences based on their nucleotide sequence, it is also possible to remove them based solely on their length using an offset. Using an offset can be useful if the sequence has many uncalled bases in the primer region, which could otherwise hinder matches between the target sequence defined in the primer table and the sequence read.

Depending on the sequence representation chosen, different algorithms are used for the removal of the subsequences described above, the dereplication of the dataset, and, if paired-end reads are used, the assembly of forward- and reverse-reads. In the OTU variant of the workflow, PANDAseq [19] is used for assembly and subsequence removal applying probabilistic error correction to assemble overlapping forward- and reverse-reads. After assembly and sequence trimming, PANDAseq will remove sequences that do not meet a minimal or maximal length threshold, have an assembly quality score below a user-defined threshold and sequences whose forward and reverse read do not have a sufficiently long overlap. The thresholds for each of these procedures can be adjusted in the configuration file. If the reads are single-end, the subsequences (poly-N, barcode, and the primer) are removed, followed by the removal of sequences that do not meet a minimal or maximal length threshold as defined in the configuration file.

The CD-HIT-EST algorithm [20] dereplicates sequences if they are either identical or if a sequence is a subsequence of another sequence. Beginning with the longest sequence of the dataset as the first representative sequence, it iterates through the dataset in order of decreasing sequence lengths, comparing at each iteration the current query sequence to all representative sequences. If the sequence identity threshold defined in the configuration file is met for a representative sequence, the counter of the representative sequence is increased by one. If the threshold could not be met for any of the existing representative sequences, the query sequence is added to the pool of representative sequences. Subsequently, the output of CD-HIT is used to count the number of sequences represented by each cluster, followed by sorting the representative sequences in descending order according to the cluster size and addition of a specific header to each sequence, as required by the VSEARCH chimera detection algorithm.

In the ASV variant of the workflow, cutadapt [21] is used to remove the subsequences defined in the primer file. After subsequence removal, DADA2 [9] is used to dereplicate the dataset and to generate ASVs using a denoising algorithm. The denoising algorithm uses a model of Illumina sequencing errors. Based on the composition, quality, and abundance of a sequence, the algorithm infers if a sequence was produced by a different sequence given the error model. After denoising, the forward and reverse reads are assembled. The assembly algorithm assumes that most substitution errors have been removed. Based on this assumption, only exactly overlapping sequences are assembled. The assembled ASVs are saved as FASTA files for downstream analysis.

#### Chimera detection

Natrix uses the VSEARCH uchime3 denovo algorithm to detect chimeric sequences. VSEARCH is an open-source alternative to the USEARCH toolkit, aimed at functionally replicating the algorithms used by USEARCH for which the source code is not openly available and that are often only described in a rudimentary manner [22]. The VSEARCH uchime3 denovo algorithm is a replication of the UCHIME2 algorithm [23] with optimized standard parameters.

#### Split-sample approach

Natrix supports both single-sample and split-sample FASTQ amplicon data. The split-sample protocol [11] aims to reduce the number of sequences that are the result of PCR or sequencing errors without using stringent abundance cutoffs, which often lead to the loss of rare but naturally occurring sequences. To achieve this, extracted DNA from a single sample is divided into two split-samples that are separately amplified and sequenced. All sequences that do not occur in both split-samples are considered as erroneous sequences and filtered out. The method is therefore based on the idea that a sequence that was created by PCR or sequencing errors does not occur in both samples. A schematic representation of the split-sample method is shown in Figure 2.

**Figure 2:**
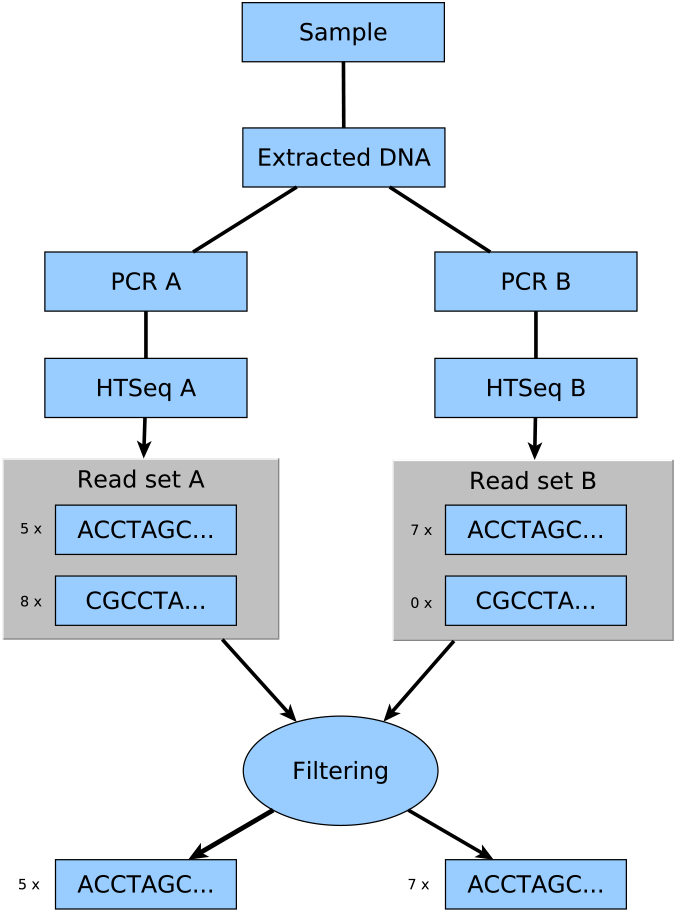
Schematic representation of the split-sample approach: Extracted DNA from a single environmental sample is split and separately amplified and sequenced. The filtering rule compares the resulting read sets between two split-samples, filtering out all sequences that do not occur in both split-samples. Image adapted from Lange et al. [11].

The initial proposal for the split-sample approach Lange et al. [11] was accompanied by the release of the R package AmpliconDuo for the statistical analysis of amplicon data produced by the aforementioned split-sample approach. It uses Fisher’s exact test to detect significantly deviating read numbers between two experimental branches A and B for each sample.

For further processing, all FASTA files are merged into a single data frame. Therefore, either an abundance cutoff value is used that can be specified in the configuration file to remove all sequences that have abundances less or equal the specified cut-off value or AmpliconDuo is applied on split-samples. The results of the discordance calculations from AmpliconDuo are plotted for visualization purposes and written to an Rdata file for further analysis. The retained and filtered sequences are subsequently exported as comma separate tables and FASTA files.

#### OTU picking

In the OTU variant of the workflow, the swarm clustering algorithm is used. Swarm clusters sequences into OTUs using an iterative approach with a local threshold: it creates a pool of amplicons from the input file and an empty OTU. Subsequently, it will remove the first amplicon from the pool, which will become the OTU seed. All amplicons left in the pool that differ in their nucleotide composition from the initial seed by a user given threshold (the default threshold used is 1 nucleotide) are removed from the pool and added to the OTU as subseeds. In the next iteration, each amplicon having at most a difference as high as the threshold to any of the subseeds is then removed from the pool and added to the OTU. This iterative process will continue until there are no amplicons left in the pool with a nucleotide difference of at most the threshold to any of the subseeds added in the previous iteration to the OTU, leading to the closure of the OTU and the opening of a new one. This approach to OTU generation circumvents two sources of OTU variability that are inherent to greedy clustering algorithms: the input order dependency, in which the first amplicon in a FASTA file will become the centroid of an OTU and the use of a global threshold, recruiting all amplicons that have fewer differences to the centroid than a user-defined threshold. The sequence of the amplicon at the center of each OTU tree is used in subsequent analysis steps as the representative sequence of the corresponding OTU.

#### Taxonomic assignment

The assignment of taxonomic information to sequence representatives is an important part of the processing of environmental amplicon data to describe and compare the communities. To find sequences that are similar to the representative sequence, the BLAST algorithm [24] is used to search for similar sequences in the SILVA and NCBI databases. The SILVA database contains aligned rRNA sequencing data that are curated in a multi-step process. The SILVA database requires 4.9 GB disk space. While it has an extensive collection of high-quality prokaryotic rRNA sequencing data, it only contains a limited amount of microbial eukaryotic sequencing data. In this case, the NCBI nucleotide (nt) database can be used, which requires 83 GB free disk space. If the database is not locally available, the required files will automatically be downloaded and the database will be built. The nucleotide-nucleotide BLAST (BLASTn) variation of the BLAST algorithm is used to assign a taxonomy for each query sequence. The best or several blast hits, as defined in the configuration file, can be used to assign a taxonomic lineage. For SILVA, with better coverage of bacterial rRNA sequences, the best hit is used to obtain the taxonomic lineage for each sequence. When the NCBI database is used, several BLAST hits should be used to automatically determine the best taxonomic assignment considering the score, length of the alignment, and resolution of the taxonomic path, which is not standardized to the same ranks as in the SILVA database. All BLAST hits with taxonomic lineage are additionally exported to a file for manual inspection, if necessary. The outputs are merged into a single comma-separated table, containing for each representative sequence the sequence identification number, the nucleotide sequence, the abundance of the sequence in each sample, and the information from the best result of the BLAST search including the taxonomic lineage.

#### Analyzed dataset

Both the OTU and the ASV variant of Natrix were applied to a subset of amplicon sequencing data taken from a study by Graupner et al. [25]. In mesocosm experiments, the effect of short-term flooding on the microbial communities in freshwater and soil was analyzed. Six AquaFlow systems were sampled at 7 time points (day1, day1-flooding1, day1-flooding2, day2, day3, day9, and day14). At each time point of sampling, water samples were collected, whereas soil samples were only collected when the AquaFlow systems were not flooded. The targeted amplicon includes an approximately 600 bp long fragment consisting of the SSU V9 region and the ITS1 (for details on study design, sampling, extraction, and PCR see [25] and [26]). Split-sample PCRs were conducted and the samples were sequenced in paired-end, rapidrun mode on the Illumina HiSeq2500 with 2×300 bp reads. Sequencing was carried out by a sequencing provider (Fasteris, Geneva). The raw sequencing data is available at NCBI, project PRJNA388564. The options chosen in the configuration file for each workflow variant are shown in additional File 4.

## 3 Discussion

### Comparison to other bioinformatics tools

In contrast to other bioinformatics tools used for amplicon analysis, Natrix is usable without extensive command-line or scripting experience and is easily extensible for more experienced users. Similar to the QIIME2 plugin system and in contrast to mothur and USEARCH, Natrix is easily extensible using Snakemake wrappers. The workflow can further be extended or modified using the rule-based Snakemake syntax and Python, R scripts, or shell commands. The ease at which Natrix can be extended is unique among amplicon analysis pipelines: while mothur requires knowledge of C++ and the mothur project itself, QIIME2 requires familiarization of the plugin system utilized by QIIME2, and USEARCH disallows the modification or extension of the source code by being closed source. Natrix can be extended using Python, R, or shell scripts with minimal adjustments. A standalone script can be incorporated into Natrix by specifying the input and output of the script in a Snakemake rule, the path to the script, and in the script itself, the only necessary adjustments are replacing the input and output file paths by a reference to Snakemake. A further advantage is the automatic parallelization of rules by Snakemake. While in other workflows the processing steps have to be run sequentially, Snakemake infers which rules are independent of each other and can be run in parallel. This reduces idle CPU time, leading to faster workflow completions. The fire-and-forget approach is another advantage, especially for large datasets and distributed computing environments: all workflow steps are configured using a single configuration file, with each entry documented. The workflow will then automatically execute all rules as specified. This is in contrast to the command-based approaches of mothur, QIIME2, and USEARCH, which will need either constant attention from the user to input new commands after the completion of the previous task or requires the ability to script a pipeline that automatically executes the commands in order. Other pipelines require the user to follow lengthy and complex tutorials while keeping track of the right input files for each step. Since the transfer of output data from one step of the workflow to the next step is handled automatically by Natrix, the user does not have to remember the file requirements for each step. This leads to a short but concise usage tutorial.

### Natrix analysis on amplicon sequences from the flooding experiment

A subset, consisting of one Illumina sequencing run, of the amplicon data from the flooding experiment was processed using the Natrix OTU and ASV variants (the parameters used are shown in additional File 4). The ASV variant of the workflow resulted in 4,770 ASVs to which taxonomic information could be assigned by BLAST, while in the OTU variant of the workflow 18,717 OTUs could be assigned taxonomically (Fig. 3). The discrepancy between the two workflow variants results from the removal of singletons and denoising by the DADA2 algorithm. On average, for each sample, 45% of the OTUs with taxonomic information contain a singleton in one of the split-samples. Since singletons are removed by DADA2, these sequences will be removed in the ASV variant of the workflow if they are not recognized as a product of a different sequence with sequencing errors. The filtering of singletons can be optionally performed on the OTU results, providing more control over the filtering steps.

**Figure 3:**
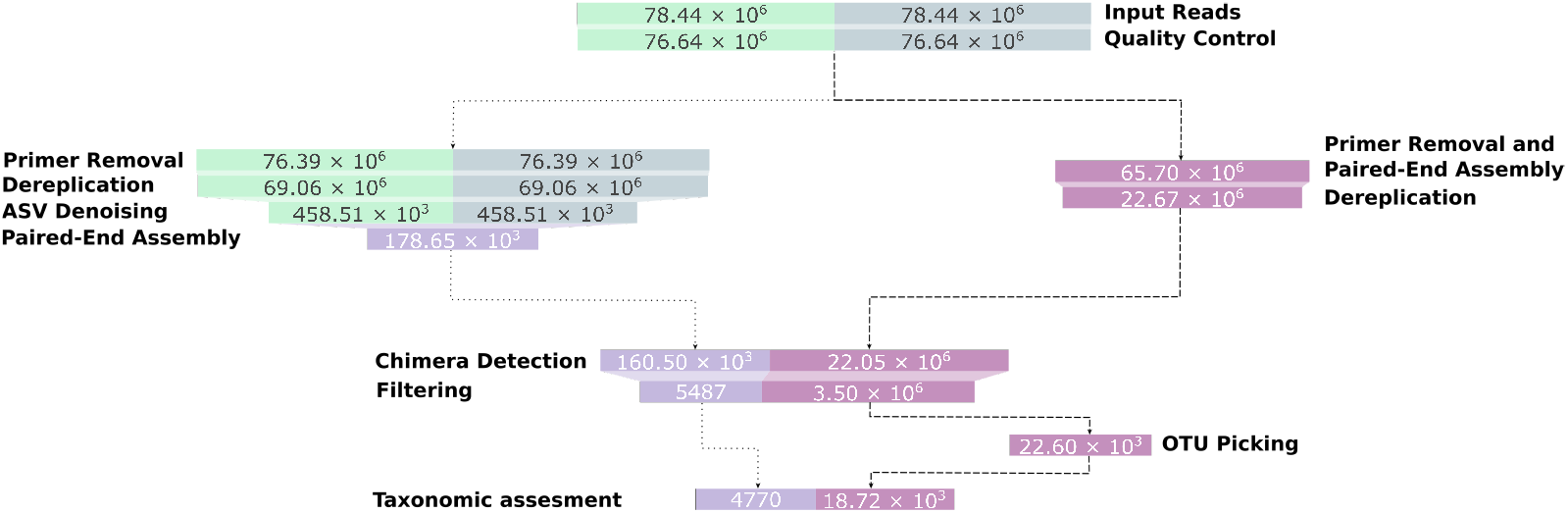
Representative sequences left after each main step of the workflow. The dotted edges represent the ASV variant of the workflow, the dashed edges represent the OTU variant. For increased visibility, the bar sizes are proportional to a logarithmic transformation of the sequence counts.

Furthermore, DADA2 identified a large number of sequences as a result of sequencing errors and added their abundance to existing ASVs. While this leads to a decreased amount of ASVs, the total read count is higher in the ASV variant of the workflow compared to the OTU variant (Fig. 4). Within the ASV variant of the workflow, DADA2 and the split-sample merging had the largest impact on the percentage of removed reads, with 41% and 43% of the total removed reads, respectively (Fig. 5). In the OTU variant of the workflow, the split-sample merging had the largest impact, with 62% of the total removed reads, followed by PANDASeq with 28% of the total removed reads.

**Figure 4:**
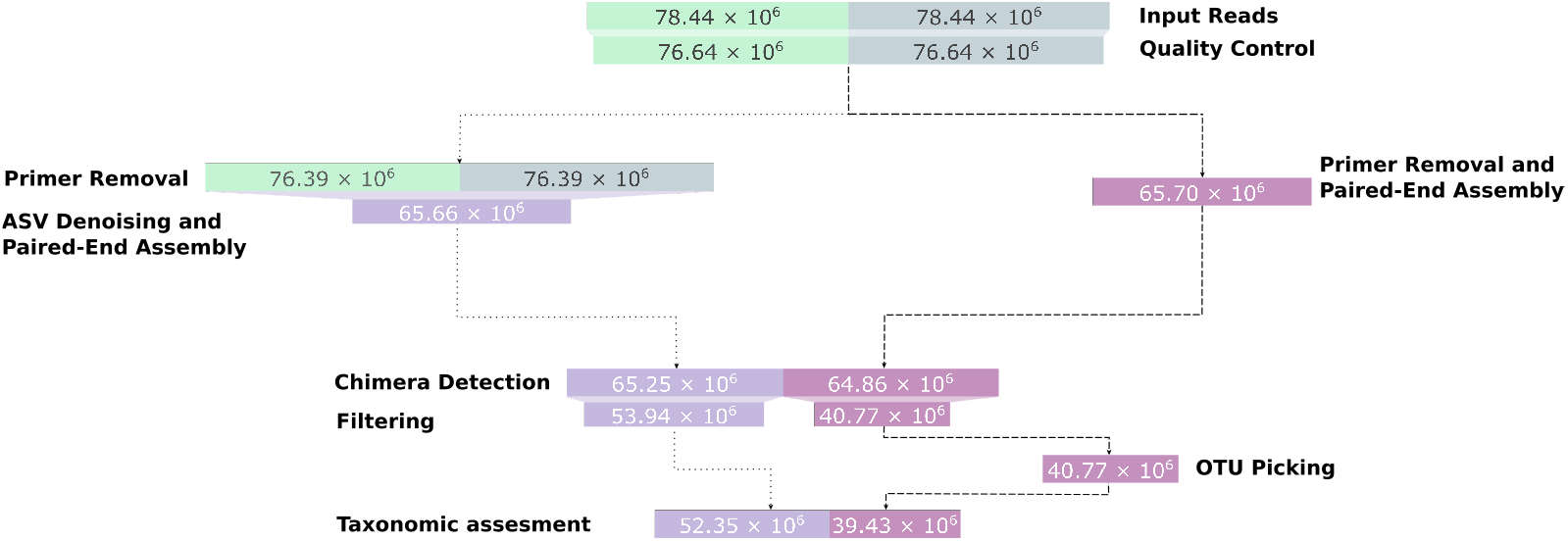
Reads left after each main step of the workflow. The dotted edges represent the ASV variant of the workflow, the dashed edges represent the OTU variant. The bar sizes are proportional to the read counts.

**Figure 5:**
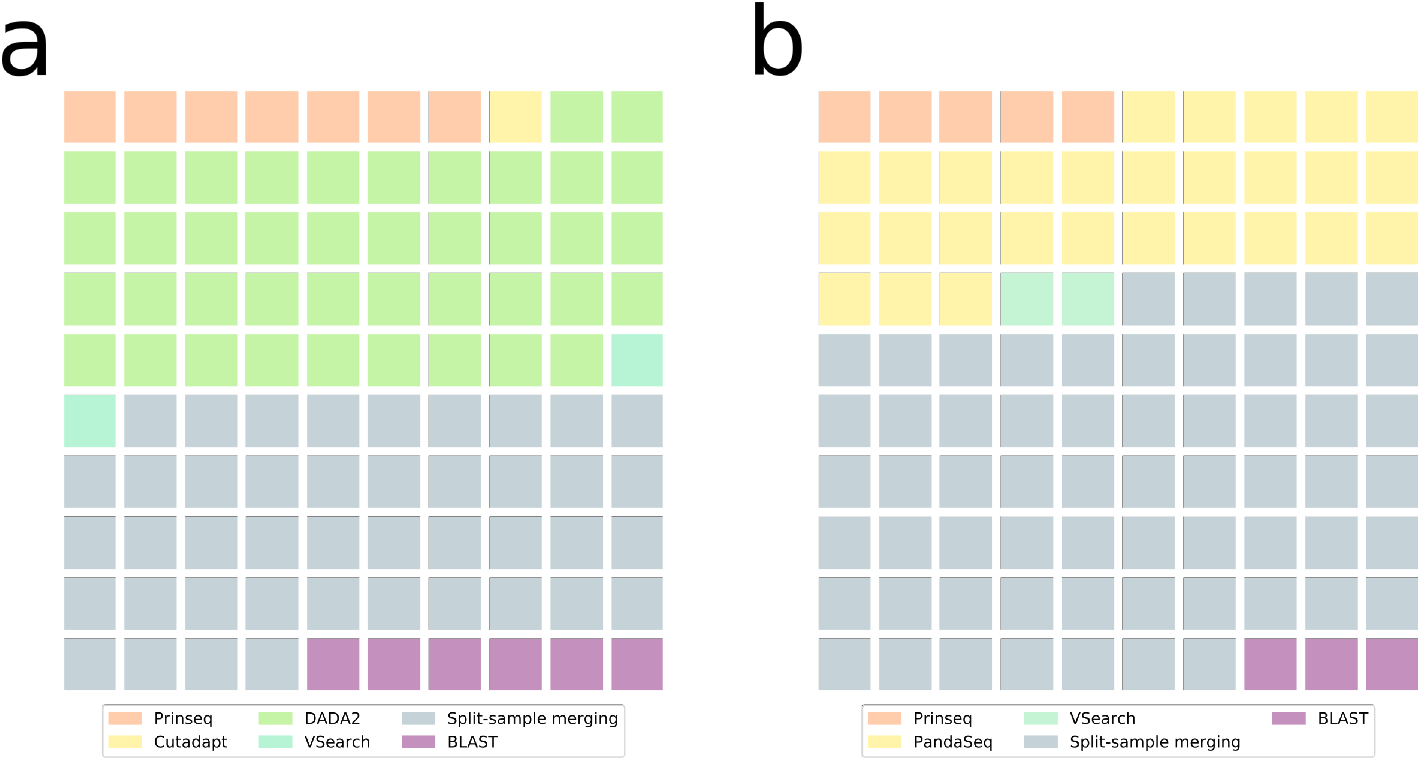
Reads removed for each main step of the ASV variant (a) and the OTU variant (b) of the workflow. Each square represents one percent of the total amount of removed reads (100 %).

From a taxonomic perspective, the results of the two workflow variants are highly similar. Both methods yield a comparable community composition (see Fig. 6 and 7). The distribution of differences between the relative abundance matrices of taxonomic groups for all samples peaks at 0.0 (Fig. 7) and the used method has no significant effect on the variation of the abundance data (adonis and anova analyses performed with the vegan [27] package). The largest difference between the workflows can be observed for Chlorophyta, followed by unclassified Fungi, Bacillariophyta and Ascomycota. When looking at the effect of flooding, the same shifts in taxonomic groups can be observed. For example, with both methods the Bacillariophyta in the soil samples become more frequent after one day of flooding (Fig. 6). This observation is consistent with the hypothesis that Bacillariophyta are indicator species for flooding events [28].

**Figure 6:**
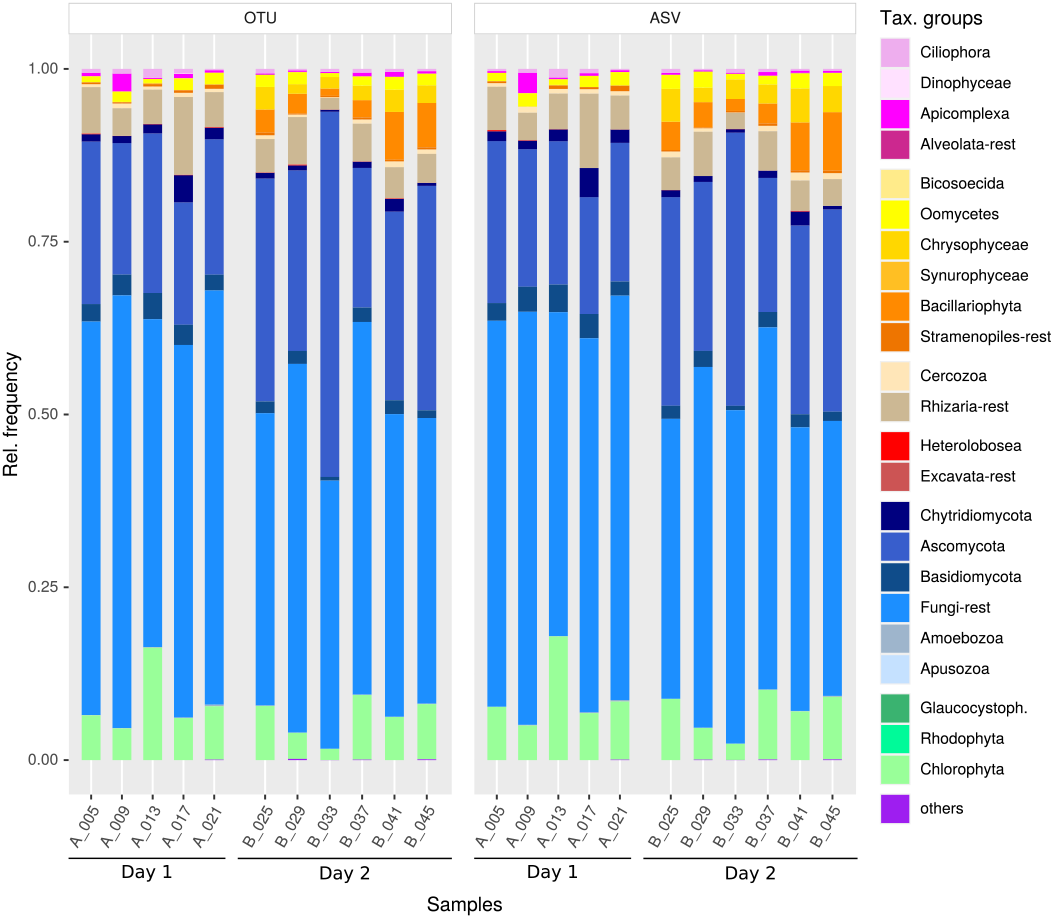
Histogram showing the relative frequencies of taxonomic groups between samples resulting from the OTU (left) and ASV (right) version of the workflow. Day 1 depicts soil samples taken before flooding (5 replicate samples; starting with A), Day 2 identifies soil samples taken one day after flooding (6 replicate samples; starting with B).

**Figure 7:**
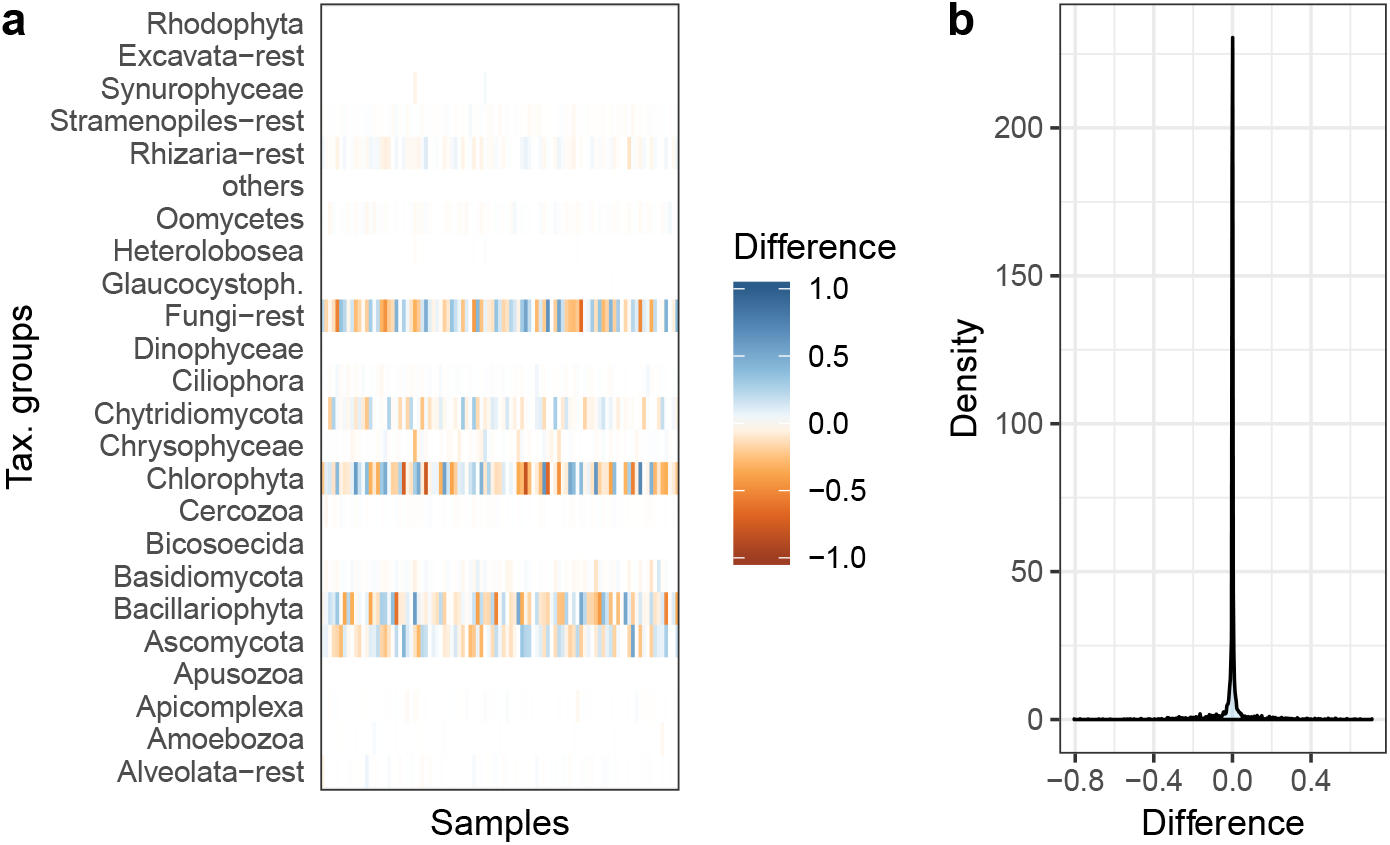
Comparison of the taxonomic composition of all 194 samples obtained for the two workflow variants. (a) depicts the differences between the relative abundance matrices of the OTU and ASV variant. Colours range between orange (difference of -1) over white (0) to blue (1). (b) depicts the distribution of the difference.

For this dataset, experimental structure, and pipeline parameters, the choice of ASV or OTU boils down to personal preference.

## 4 Conclusion

Natrix allows amplicon processing from raw Illumina reads up to the taxonomic assessment of sequence representatives with a single command. It requires no programming experience and scales seamlessly to different computing environments. All applications and algorithms incorporated into Natrix can be fine-tuned in the accompanying configuration file, with each option having a detailed description and being safeguarded by sanity checks. The rule-based Snakemake syntax allows easy modification and extension of Natrix or the incorporation of parts into other workflows. Natrix is the only amplicon processing workflow that supports the Amplicon-Duo split-sample approach, reducing the number of sequences that are the result of PCR or sequencing errors. It further supports different sequence representations in the form of OTUs or ASVs, allowing researchers to choose either of them, depending on the study type that is performed. Switching between the different sequence representations can be performed by a single entry in the configuration file, and Natrix also facilitates a comparison between them. The combination of user-friendliness, high modularity, and extensibility makes Natrix an attractive amplicon processing workflow for a wide range of potential users.

## 5 Availability and requirements

Project name: Natrix

Project home page: https://github.com/MW55/Natrix

Operating system(s): Linux

Programming environment: Snakemake, Python, R, Bash

Other requirements: Python 3.7 or higher, Conda

License: MIT

Any restrictions to use by non-academics: N / A

Natrix is free and open-source software. It is available on GitHub (https://github.com/MW55/Natrix). The GitHub page contains extensive documentation and a tutorial. Natrix depends, besides on Snakemake, on the Conda package manager. Conda can be downloaded as part of the Anaconda or the Miniconda platforms (Python 3.7). All other dependencies will be automatically installed using Conda environments and can be found in the corresponding environment.yaml files in the envs folder and the snakemake.yaml file in the root directory of the workflow.

