## Supplement for "Natrix: A Snakemake-based workflow for processing, clustering, and taxonomically assigning amplicon sequencing reads"

May 2020

### Additional files

#### Additional file 1 — Output file hierarchy

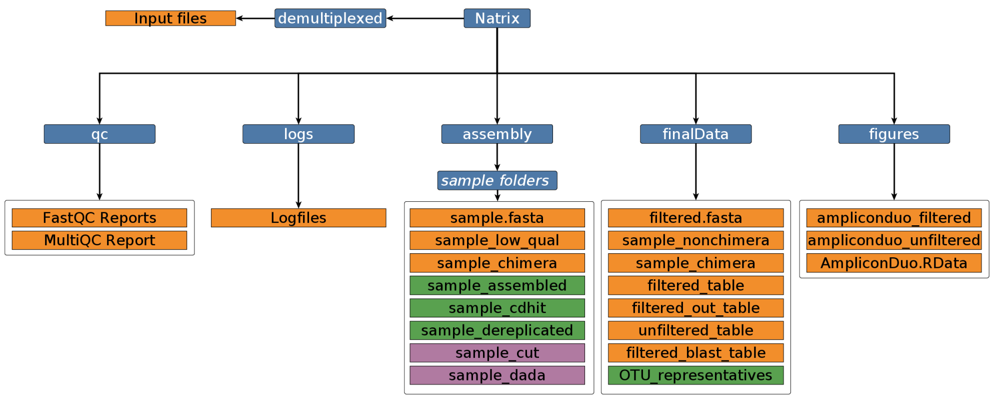


Figure 1: Output file hierarchy, blue nodes represent folders, orange nodes represent files that are created in both variants of the workflow, green nodes are files exclusive to the OTU variant and purple nodes are files exclusive to the ASV variant of the workflow..

#### Additional file 2 — Output file descriptions

| Folder | File(s) | Description |
|---|---|---|
| qc | FastQC reports | Quality reports of the FastQC application. |
| | MultiQC report | Aggregated FastQC reports in a single file. |
| logs | Logfiles | Logfiles of the different rules. |
| assembly (one folder for each sample) | sample_low_qual.fastq | Sequences of sample that did not pass the prinseq quality filtering. |

| | | |
|---|---|---|
| | sample_assembled.fastq | With PANDAseq assembled sequences. |
| | sample_CDHIT.fastq | Representative sequences of dereplication clusters. |
| | sample.fasta | FASTA file of the assembled sequences. |
| | sample.dereplicated.fasta | Dereplicated sequences of sample |
| | sample_chimera.fasta | Sequences of sample that are thought to be of chimeric origin. |
| | sample_cut.fasta | Sequences of sample without additional subsequences (primer, barcodes etc.). |
| | sample_dada.fasta | ASVs of sample that underwent denoising, dereplication and paired-end merging using the DADA2 toolkit. |
| finalData | sample.nonchimera.fasta | Sequences of sample that passed the chimera detection rule. |
| | unfiltered_table.csv | Table containing the sequences of all samples and their abundances per sample. |
| | filtered_table.csv | Table containing the sequences of all samples and their abundances per sample after filtering. |
| | filtered_out_table.csv | Table containing the sequences that did not pass the filtering rule. |
| | filtered.fasta | The sequences of the filtered_table.csv in FASTA format. |
| | filtered_blast_table.csv | Table containing the sequences of the filtered_table.csv and the taxonomic information assigned to each. |
| figures | ampliconduo_unfiltered.png | Discordance graph before the filtering. |
| | ampliconduo_filtered.png | Discordance graph after filtering. |
| | AmpliconDuo.Rdata | RData file containing the results of the AmpliconDuo statistical analysis. |

### Additional file 3 — Example configuration file with default values

For the configuration of sub-workflows and applications used by Natrix, a single configuration file is used. Optional parts of the workflow (e.g., the generation of quality reports, clustering of OTUs, or the assignment of taxonomic information) can be disabled. The configuration file is also used to define the protocols used to generate the input data, e.g., whether the data is in single-end or paired-end format and whether a split-sample approach was used during sample preparation. It also contains configuration options to adjust individual parts of the workflow depending on the requirements of the project.

An example configuration file with a description and default value for each parameter is shown in Table 2.

| **Option** | **Default** | **Description** |
|---|---|---|
| filename | project | The filename of the project folder, primertable (.csv) and config file (.yaml). |

| | | |
|---|---|---|
| primertable | project.csv | Path to the primertable. |
| units | units.tsv | Path to the sequencing unit sheet. |
| cores | 4 | Amount of cores available for the workflow. |
| multiqc | True | Initial quality check (fastqc & multiqc), currently only works for not yet assembled reads. |
| demultiplexing | False | Demultiplexing for reads that were not demultiplexed by the sequencing company (slow). |
| read_sorting | False | Read sorting for paired end reads that were not sorted by the sequencing company (slow). |
| already_assembled | False | Skipping of the quality control and read assembly steps for data that is already assembled. |
| seq_rep | OTU | How the sequences should be represented, possible values are: ”ASV”, amplicon sequence variants, created with DADA2 or ”OTU”, operational taxonomic units, created with SWARM |
| threshold | 0.9 | PANDAseq score threshold a sequence must meet to be kept in the output. |
| minoverlap | 15 | Sets the minimum overlap between forward and reverse reads. |
| minqual | 1 | Minimal quality score for bases in an assembled read to be accepted by PANDAseq. |
| minlen | 100 | The minimal length of a sequence after primer removal to be accepted by PANDAseq. |
| maxlen | 600 | The maximal length of a sequence after primer removal to be accepted by PANDAseq. |
| primer_offset | False | Using PANDAseq to remove primer sequences by length offset instead of sequence identity. |
| mq | 25 | Minimum quality sequence check (prinseq), filtering of sequences according to the PHRED quality score before the assembly. |
| barcode_removed | True | Boolean that indicates if the sequence is free of barcodes. |
| all_primer | True | Boolean that indicates if the sequence is free of any kind of additional subsequences (primer, barcodes etc.). |

| | | |
|---|---|---|
| clustering | 1.0 | Percent identity for cdhit (dereplication) (1 = 100%), if cdhit is solely to be used for dereplication (recommended), keep the default value. |
| length_overlap | 0.0 | Length difference cutoff, default 0.0 if set to 0.9, the shorter sequences need to be at least 90% length of the representative of the cluster. |
| representative | longest | Which sequence to use as a representative sequence per CDHIT cluster. longest = the longest sequence of the corresponding cluster, most_common = the most common sequence of the corresponding cluster. |
| beta | 8.0 | Weight of a ”no” vote for the VSEARCH chimera detection algorithm. |
| pseudo_count | 1.2 | Pseudo - count prior on number of “no” votes. |
| abskew | 16 | Minimum abundance skew, definied by (min(abund.(paren1), abund.(paren2))) / abund.(child). |
| filter_method | not_split | If the split sample approach was used (split_sample) or not (not_split). |
| cutoff | 3 | An additional abundance filter if the split sample approach was not used. |
| ampli_corr | fdr | Specifies the correction method for Fisher’s exact test. |
| save_format | png | File format for the frequency-frequency plot. |
| plot_AmpDuo | True | If the frequency-frequency plot should be saved. |
| paired_End | True | The format of the sequencing data, TRUE if the reads are in paired-end format. |
| name_ext | R1 | The identifier for the forward read (for the reverse read the 1 is switched with 2, if the data is in paired-end format), has to be included at the end of the file name, before the file format identifier (including for single end files). |
| swarm | True | Boolean to indicate the use of the SWARM clustering algorithm to create operational taxonomic units (OTUs) from the data. |
| blast | True | Boolean to indicate the use of the BLAST clustering algorithm to assign taxonomic information to the OTUs. |

| | | |
|---|---|---|
| database | SILVA | Database against which the BLAST should be carried out, at the moment ”NCBI” and ”SILVA” are supported. |
| drop_tax_classes | '.*unclassified Bacteria.*, .*uncultured.*bacterium.*' | Given a comma-separated list, drops undesired classes either by id, by name or using regex. |
| db_path | database/silva.db | Path to the database file against which the BLAST should be carried out, at the moment only the SILVA and NCBI databases will be automatically downloaded, other databases have to be downloaded and configurated manually. |
| max_target_seqs | 1 | Number of blast hits that are saved per sequence / OTU. |
| ident | 90.0 | Minimal identity overlap between target and query sequence. |
| evalue | 1e-51 | Highest accepted evalue. |
| out6 | ”6 qseqid qlen length pident mismatch qstart qend sstart send gaps evalue stitle” | Additional BLAST information to be saved. |

Table 2: Example configuration file with default values.

### Additional file 4 — Configuration parameters used for the flooding experiment.

| **Option** | **OTU Workflow** | **ASV Workflow** |
|---|---|---|
| filename | JAU-4_otu | JAU-4_asv |
| primertable | JAU-4_otu.csv | JAU-4_asv.csv |
| units | units.tsv | units.tsv |
| cores | 20 | 20 |
| multiqc | False | False |
| demultiplexing | False | False |
| read_sorting | False | False |
| already_assembled | False | False |
| seq_rep | OTU | ASV |
| threshold | 0.9 | 0.9 |
| minoverlap | 15 | 15 |
| minqual | 1 | 1 |
| minlen | 100 | 100 |
| maxlen | 600 | 600 |
| primer_offset | False | False |
| mq | 30 | 30 |
| barcode_removed | True | True |
| all_primer | False | False |
| clustering | 1.0 | 1.0 |
| length_overlap | 0.0 | 0.0 |
| representative | most_common | most_common |
| beta | 8.0 | 8.0 |

| | | |
|---|---|---|
| pseudo_count | 1.2 | 1.2 |
| abskew | 16 | 16 |
| filter_method | split_sample | split_sample |
| cutoff | 3 | 3 |
| ampli_corr | fdr | fdr |
| save_format | png | png |
| plot_AmpDuo | True | True |
| paired_End | True | True |
| name_ext | R1 | R1 |
| swarm | True | False |
| blast | True | True |
| database | NCBI | NCBI |
| drop_tax_classes | ” | ” |
| db_path | database/ncbi/nt | database/ncbi/nt |
| max_target_seqs | 10 | 10 |
| ident | 90.0 | 90.0 |
| evalue | 1e-51 | 1e-51 |
| out6 | ”6 qseqid qlen length pident mismatch qstart qend sstart send gaps evalue stitle” | ”6 qseqid qlen length pident mismatch qstart qend sstart send gaps evalue stitle” |

Table 3: Configuration parameters used for the flooding experiment.

### Additional file 5 — Representative sequence and read counts of the flooding experiment.

ODS file containing the read counts and the amount of representative sequences left after each main step of the workflow.
